# Cohort profile: The LipidCardio Study - Role of Lipoproteins in Cardiovascular Disease

**DOI:** 10.1101/669150

**Authors:** Maximilian König, Samita Joshi, David M. Leistner, Ulf Landmesser, David Sinning, Elisabeth Steinhagen-Thiessen, Ilja Demuth

## Abstract

**Purpose:** The LipidCardio Study was established for in-depth analyses of cardiovascular risk factors, providing well-defined cardiovascular and metabolic phenotypes. Particularly the role of lipoproteins in the pathobiological process and treatment of cardiovascular disease will be a main focus.

**Participants:** 1.005 individuals aged 21 years and older undergoing cardiac catheterization during 17 months at a tertiary academic cardiology center were enrolled. The baseline data set contains detailed phenotyping, broad biochemical parameters, genetic data, but also standardized personal and family history, a screening test for cognitive impairment, pulse wave analysis and measurements of hand grip strength, amongst others. Blood samples were stored in a biobank for future analyses.

**Findings to date:** The mean age of the participants at enrolment was 70.9±11.1 years (70% male). Coronary angiography provided evidence of obstructive coronary artery disease (CAD) in 69.9% of participants. Those with evidence of CAD were significantly more likely to be male, inactive, diabetic and with a family history of cardiovascular disease than participants without CAD.

20% of patients had lipoprotein(a) [Lp(a)] concentrations above 106.9 nmol/L (fifth quintile). These patients had significantly increased odds of obstructive CAD compared to participants in quintiles 1-4 (OR 1.70, 95% CI 1.17 to 2.48, p=0.005). There was reasonable evidence that with increasing severity of CAD the odds of having elevated Lp(a) increased. We were able to replicate the established strong association between specified single nucleotide polymorphisms (SNPs) in the *LPA gene* (rs10455872, rs3798220 and rs186696265) and the *APOE gene* (rs7412), and the concentration of Lp(a), validating our phenotype database and biobank.

**Future plans:** Mortality information will be obtained in two-year intervals. Follow-up phone interviews will be conducted at 3, and 6 years after enrolment. We seek to cooperate with other researchers in the field, e.g. by sharing data and biobank samples.

**Registration:** not applicable, purely observational study

## Introduction

Ischemic heart disease accounts for > 20% of all deaths in Europe. [1] Although age-adjusted mortality rates from coronary artery disease (CAD) have declined substantially in the past decades throughout Europe and the US, the incidence and prevalence of CAD remain high. [2, 3] Accordingly, 27% of all disability-adjusted life years (DALYs) in Germany are due to coronary artery disease.[1]

It is well established that traditional modifiable risk factors for CAD (hypertension, hypercholesterolemia, sedentary lifestyle, obesity smoking, and diabetes) predict the risk of *developing* CAD, and targeting them decreases cardiovascular risk in a dose-dependent manner.[4] However, among patients with prevalent CAD, it has been shown that there is substantial variation in estimated 10-year risk of outcomes, with a large proportion of individuals with a 10-year risk <10%, but also a considerable proportion with very high residual risk despite optimally controlled risk factors.[5] Therefore, the identification and consideration of additional modifiable risk factors to reduce (residual) cardiovascular risk is crucial, and novel biomarkers, and assessment of the individual genetic profile bear the potential of improving risk assessment for primary and secondary prevention. Since patients with prevalent CAD are a very heterogeneous group, discrimination is likely to be advantageous over a one fits all disease management strategy.

Dyslipidemia is a major cause of atherosclerotic cardiovascular disease. In addition to low density lipoprotein cholesterol (LDL-C), lipoprotein (a) [Lp(a)] has emerged as an independent risk factor for CAD. [6] Whereas the causal role of LDL-C is well established, Lp(a) is less well understood and still a subject of controversy. [6, 7] The underlying physiological function of this lipoprotein is still unknown. Several studies have indicated that serum/plasma Lp(a) is largely genetically determined, and there is strong suggestion that Lp(a) is causally associated with CAD.[8] Prospective observational studies have shown that Lp(a) was an independent risk factor for major adverse cardiovascular events (MACEs), i.e. the progression of CAD in patients with stable CAD who received optimal medical therapy, and that Lp(a) levels were positively associated with coronary lesion severity as well.[7, 9]

Screening for elevated Lp(a) presently is recommended in those at intermediate or high CV risk, in particular as an additional risk indicator.[7, 10, 11] Moreover, Lp(a) has been suggested as a potential therapeutic target.[12] To note, an antisense oligonucleotide specific to apo(a) has been recently shown to lead to dose-dependent reductions in average Lp(a) levels of >80%.[13]

Overall, substantial progress has been made in the field of CAD in the last decades. However, further improvements will presuppose careful characterization of different phenotypes and subsequent joint application of genetics, epigenetics and metabolomics. For this purpose we established the LipidCardio Study in 2016. We sought to implement detailed phenotyping and follow-up of patients with different severity of CAD and also without obstructive CAD to examine factors associated with the pathobiological process and progression of CVD. There will be a strong focus on lipoproteins and genetics. In particular, we seek to gain further insight into the role of Lp(a) in cardiovascular pathogenesis and to investigate the implications of a screening for Lp(a) in a high risk cohort of patients undergoing cardiac catheterization and coronary angiography. Combining phenotype factors with genetic factors, we desire to elaborate the particular cardiovascular risk profile of patients with high Lp(a).

As another integral part of this study we collected ample blood specimens to hold high-quality, well-characterized biomaterials along with their clinical, genetic, and demographic information for future biomedical research projects in store in a high-quality biobank.

### Cohort description

Patients aged 18 years and older undergoing cardiac catheterization at a single large academic center (Department of Cardiology, Campus Benjamin Franklin, Charité-Universitätsmedizin Berlin), except those with troponin-positive acute coronary syndromes (ACS), were eligible for inclusion. Participation rate was particularly high (>95%). Between October 2016 and March 2018 1,005 consecutive patients were enrolled. All participants gave written informed consent at the time of enrolment. Patients unable to provide informed consent were excluded from the study. The study was approved by the ethics committee at Charité-Universitätsmedizin Berlin (approval number: EA1/135/16).

The main study variables are shown in **Supplementary Table 1**. A comprehensive data collection was performed, with a strong focus on cardiovascular health. Blood specimens were collected at the time of cardiac catheterization (after administration of heparin), either as arterial blood sample directly from the radial or femoral artery sheath or as a venous blood sample from a peripheral intravenous access. At enrolment, patients were interviewed to collect information on demographic characteristics, over-the-counter (OTC) medication usage, health behaviors (alcohol/drug use, smoking, physical activity), a detailed standardized family history was obtained, and a screening test for cognitive impairment (the mini-mental state examination, (MMSE) was performed. Information on regular prescription medications were obtained from the patients’ medication plan. In addition, electronic medical records were reviewed by study personnel to obtain previous diagnoses of chronic conditions, as well as to document previous angiographic findings and coronary revascularization history.

Portions of the blood specimens were used to determine a basic panel of laboratory tests essential to this study. Lp(a) was determined using a turbidimetric assay (Roche Diagnostics GmbH, Mannheim). Lp(a) is given in nmol/L, correctly reflecting the number of Lp(a) particles.[14] A range of other parameters were taken over from the hospital information system (HIS), provided that they were ordered as part of the clinical routine. Importantly, all tests were done in one central, accredited laboratory.

#### Specimens

EDTA blood samples were frozen immediately following collection at −80°C until DNA was isolated using the *sbeadex livestock kit* (LGC Genomics GmbH, Germany). Selected SNPs were genotyped using KASP chemistry (LGC, Hoddesdon, UK). Additional blood samples were drawn and stored in the *Central Biomaterial Bank Charité* (ZeBanC) [15], providing the opportunity for additional measurements at a later date.

#### Coronary angiography

Cardiac catheterization and coronary angiography were performed according to the standard protocols of the Interventional Cardiology Unit and by discretion of the interventional cardiologist. The interventional cardiologist routinely documented diagnostic findings. Comprehensive angiographic results at the time of enrolment were recorded in the study database.

#### Echocardiography, blood pressure measurement, pulse wave analysis

Echocardiography was performed as a part of the clinical routine, and selected parameters were taken over into the study database.

Blood pressure was measured once on each arm, in a sitting position and at rest with a boso medicus uno^®^ device with an adequate cuff size.

Central blood pressure, pulse wave reflection, and pulse wave velocity were measured with the Mobil-O-Graph^®^ PWA device, according to the operating instructions of the manufacturer.

#### Other measurements

Waist circumference was measured at the midpoint between the lower margin of the least palpable rib and the top of the iliac crest. Hip circumference was measured around the widest portion of the buttocks, with the tape parallel to the floor. For both measurements, the subjects were standing and wore little clothing. [16]

Hand grip strength was measured with a Smedley Dynamometer (Saehan, Type SH5002).

#### Questionnaires

The Seattle Angina Questionnaire, and the Rapid Assessment of Physical Activity (RAPA) questionnaire were administered. Folstein’s Mini-Mental State Examination (MMSE) for screening of cognitive impairment was administered in face-to-face interviews by trained study personnel.

### Planned follow-up measures

Mortality information will be obtained from the obligatory registry office in two-year intervals. Follow-up phone interviews will be conducted by trained study personnel, and the calls are planned to be made at 3, and 6 years after initial enrolment. Adverse CVD events, including non-fatal MI, acute coronary syndromes, heart failure, hospitalizations, cardiac procedures (e.g., revascularization), strokes and peripheral arterial disease events will be recorded. Additional information will be collected on follow-up coronary angiogram data, development of comorbidities (hypertension, diabetes, cardiac arrhythmia, valvular heart disease, obstructive sleep apnoea and cancer) and changes in medications.

### Findings to date

#### The cohort

Baseline characteristics of the 1,005 patients are shown in Tables 1 and 2. The mean age at enrolment was 70.9±11.1 years. Approximately 70% of the participants were male, 97.6 were of white Caucasian ancestry. Coronary angiography provided evidence of obstructive coronary artery disease (CAD, defined as >50% luminal narrowing in a major epicardial vessel) in 69.9% of participants at enrolment, 11.2% had non-obstructive CAD and 18.8% had normal coronary angiograms without evidence of atherosclerosis (no apparent CAD). In 509 patients (50.6%) obstructive CAD had been previously known, and angiography was performed for evaluation of disease progression or planned intervention of residual stenosis.

**Table 1.** Baseline characteristics of the LipidCardio Study, total and according to presence or absence of elevated Lipoprotein(a) (≥ 107 nmol/L).

|  | **N** | **Total** | **Lp(a) < 107**<br>**nmol/L\*** | **Lp(a) $\geq 107$**<br>**nmol/L\*** |
| --- | --- | --- | --- | --- |
| Sex, male | 1005 | 701(69.8) | 554(71.5) | 126(65.3) |
| Age, years | 1005 | 70.9±11.1 | 70.7±11.4 | 71.7±9.9 |
| Caucasian ancestry | 1005 | 981(97.7) | 756(97.6) | 188(97.4) |
| Previous diagnosis of CAD | 1005 | 509(50.7) | 374(48.3) | 115(59.6) |
| Previous bypass surgery | 1005 | 77(7.7) | 51(6.6) | 24(12.4) |
| Previous myocardial<br>infarction | 1005 | 310(30.9) | 227(29.3) | 61(36.8) |
| Total obstructive CAD | 1005 | 701(69.9) | 526(67.9) | 151(78.2) |
| - 1 vessel | 1005 | 156(15.5) | 122(15.7) | 27(14.0) |
| - 2 vessel | 1005 | 234(23.3) | 174(22.5) | 53(27.5) |
| - 3 vessel | 1005 | 311(31.0) | 230(29.7) | 71(36.8) |
| Non-obstructive CAD | 1005 | 113(11.2) | 94(12.1) | 14(7.3) |
| No apparent CAD | 1005 | 170(16.9) | 155(20) | 28(14.5) |
| Aortic valve stenosis | 1005 | 100(12.2) | 71(11.2) | 26(16.4) |
| Previous stroke | 1005 | 98(9.8) | 71(9.2) | 24(12.4) |
| Atrial fibrillation | 1005 | 273(27.2) | 214(27.6) | 49(25.4) |
| Diabetes mellitus Type 2 | 1005 | 270(26.9) | 220(28.4) | 43(22.3) |
| Hypertension | 1005 | 813(81.0) | 627(80.9) | 156(80.8) |
| Dyslipidaemia | 1005 | 598(59.6) | 455(58.7) | 121(62.7) |
| Cancer | 1005 | 181(18.0) | 136(17.6) | 42(21.8) |
| PAD | 1005 | 91(9.1) | 64(8.3) | 23(11.9) |
| CKD | 1005 | 164(16.3) | 121(15.6) | 35(18.1) |
| COPD | 1005 | 109(10.8) | 87(11.2) | 17(8.8) |
| LDL-C (mg/dL) | 964 | 99.1±40.6 | 97.1±39.8 | 106.4±41.9 |
| ApoA1 (g/L) | 955 | 1.28±0.28 | 1.27±0.28 | 1.32±0.27 |
| ApoB (g/L) | 956 | 0.85±0.27 | 0.83±0.27 | 0.93±0.28 |
| WC (cm) | 852 | 101.7±14.1 | 102.1±14.4 | 100.6±13.6 |
| Systolic blood pressure<br>(mmHg) | 857 | 134.0±21.3 | 134.0±21.5 | 134.3±21.1 |
| Statin therapy | 1005 | 608(60.5) | 459(59.2) | 129(66.8) |
| eGFR CKD-EPI formula<br>(ml/min/1.73m<sup>2</sup>) | 972 | 70.0±19.7 | 70.4±19.7 | 69.2±19.6 |
| eGFR< 60ml/min/1.73m<sup>2</sup> | 972 | 290(29.8) | 218(29.3) | 59(30.9) |

\*Lp(a) was available only for 968 participants
Notes: Values are mean±SD and n(%) unless stated otherwise. Lp(a), Lipoprotein(a), eGFR, estimated glomerular filtration rate, CKD-EPI, Chronic Kidney Disease-Epidemiology collaboration, CAD = Coronary artery disease, TIA = Transient ischemic attack, AF = Atrial fibrillation, DM 2 = type 2 diabetes mellitus, PAD = peripheral arterial disease, CKD = Chronic kidney disease, COPD = Chronic obstructive pulmonary disease, WC, waist circumference.

**Table 2.** Baseline characteristics by **coronary artery disease** (CAD) status at baseline.

|  | **N** | **No apparent**<br>**CAD** | **Obstructive**<br>**CAD** |  | **Non-**<br>**obstructive**<br>**CAD** |
| --- | --- | --- | --- | --- | --- |
|  |  |  | **previous** | **newly diagnosed** |  |
|  | 1,003* | 187(18.6%) | 509(50.8%) | 200(19.9%) | 107(10.7%) |
| Age (years) | 1003 | 66.2±12.9 | 72.0±10.7 | 71.9±9.8 | 72.3±9.2 |
| Sex (female) | 1003 | 107(57.2) | 95(18.7) | 58(29.0) | 42(39.3) |
| Diabetes mellitus 2 | 1003 | 38(20.3) | 157(30.8) | 52(26.0) | 23(21.5) |
| Dyslipidemia | 1003 | 80(42.8) | 354(69.8) | 107(53.5) | 56(52.3) |
| Smoking (current) | 898 | 33(20.8) | 82(17.9) | 40(22.2) | 15(15) |
| Smoking (former) | 727 | 61(48.4) | 247(65.7) | 81(57.9) | 44(51.8) |
| LDL-C (mg/dl) | 962 | 115.00±37.60 | 85.20±35.13 | 117.75±44.40 | 104.38±37.53 |
| HDL-C (mg/dl) | 959 | 56.76±17.76 | 47.84±15.34 | 50.91±15.19 | 55.82±18.30 |
| Triglycerides<br>(mg/dl) | 714 | 112(88-156) | 120(89-174) | 124(89-185) | 119(89-158) |
| Lp(a) (nmol/L) | 968 | 17.4(7.2-60.3) | 20.5(5.0-99.8) | 22.3(7.2-71.4) | 15.30(5.0-44.4) |
| Lp(a) >106.9 nmol/l | 968 | 28(15.5) | 115(23.5) | 37(19.0) | 13(12.6) |
| Glucose (mg/dl) | 753 | 115.0±37.3 | 123.1±47.1 | 125.7±47.7 | 121.9±44.6 |
| HbA1c (%) | 981 | 5.70±0.70 | 6.00±0.84 | 5.99±0.89 | 5.78±0.81 |
| ApoB (g/L) | 956 | 0.93±0.24 | 0.77±0.25 | 0.95±0.30 | 0.89±0.27 |
| CKDcalc | 970 | 36(20.3) | 175(34.9) | 55(29.1) | 23(22.3) |
| eGFR<sub>CKD-EPI</sub><br>(ml/min/1.73m<sup>2</sup>) | 970 | 75.4±20.4 | 68.3±19.4 | 70.1±21.0 | 69.5±15.1 |
| ApoA1 (g/L) | 955 | 1.39±0.31 | 1.23±0.26 | 1.30±0.25 | 1.33±0.27 |
| BMI (kg/m<sup>2</sup>) | 911 | 27.89±5.05 | 27.83±4.70 | 27.58±5.03 | 27.75±5.40 |
| CRP (mg/l) | 703 | 2.15(1.1-9.1) | 2.0(0.9-6.9) | 2.8(1.1-7.6) | 2.30(0.8-7.0) |
| SBP (mmHg) | 855 | 131.96±20.40 | 133.3±21.5 | 138.3±20.8 | 132.48±20.37 |
| DBP (mmHg) | 855 | 80.6±13.0 | 76.5±13.8 | 80.5±14.1 | 76.1±10.9 |

\*N=2 patients' CAD status (0.2%) was "other". Notes: Values are mean ± SD and n(%) or median (25<sup>th</sup>-75<sup>th</sup> percentile) unless stated otherwise. eGFR = estimated glomerular filtration rate, CKD-EPI, Chronic Kidney Disease-Epidemiology collaboration, CAD = Coronary artery disease, TIA = Transient ischemic attack, AF = Atrial fibrillation, DM 2 = type 2 diabetes mellitus, PAD = peripheral arterial disease, CKD = Chronic kidney disease, COPD = Chronic obstructive pulmonary disease, BMI = body mass index, CRP = C reactive protein, SBP/DBP systolic/diastolic blood pressure, ApoA1 apoprotein A1, ApoB, apoprotein B, LDL-C, low density lipoprotein cholesterol, HDL-C, high-density lipoprotein cholesterol

Those with evidence of obstructive CAD were significantly more likely to have a positive family history of cardiovascular disease (odds ratio (OR) 1.48, 95% confidence interval (CI) 1.06 to 2.06, p=0.019), compared to participants with no apparent CAD. Likewise, participants with obstructive CAD were significantly more likely to be physically inactive (OR 1.50, 95% CI 1.09 to 2.16, p=0.014).

### Lipoprotein(a) – clinical and genetic associations

The median serum concentration of Lp(a) at baseline was 19.3 nmol/L (interquartile range [IQR] <7.0 to 77.1 nmol/L). Lp(a) levels were positively associated with age among women, whereas in men Lp(a) levels were consistent across the age spectrum. Referring to this, whereas overall there was no evidence of a significant sex difference in Lp(a) levels, in patients aged over 60 years the median Lp(a) concentration was significantly higher in women (26.4; IQR 8.2 to 109.6) than in men (18.2; IQR <7.0 to 71.4, p=0.038).

Accounting for the right-skewed distribution of Lp(a) in the general population and also in cardiovascular high-risk populations, it has become current practice to examine the distribution of Lp(a) by quintiles, particularly individuals in the fifth quintile being at increased risk of CV disease. Also, the commonly referenced threshold of 100-125 nmol/L (equals 50 mg/dL), usually corresponds approximately to the cut-off point defining the fifth quintile. [6, 17] Of total 968 patients with available Lp(a) measurements 193 patients (19.9%) had Lp(a) values above 106.9 nmol/L (fifth quintile).

Participants in the highest quintile of Lp(a) (≥ 107 nmol/L) had significantly increased odds of obstructive CAD compared to participants in the lowest quintile (OR 1.58, 1.02 to 2.45, p=0.039) or in quintiles 1-4 (OR 1.7, 95% CI 1.17 to 2.48, p=0.005), respectively. Moreover, there was reasonable evidence that with increasing severity of CAD (non obstructive CAD, 1-vessel, 2-vessel, and 3-vessel obstructive CAD) the odds of having a significantly elevated Lp(a) increased (test for trend p=0.015).

#### Genetic associations

In order to assess if our cohort data are suitable for genetic association analyses, we genotyped and examined *LPA*-SNPs rs10455872, rs3798220 and rs186696265, which have been previously reported to be strongly associated with Lp(a) serum levels. [18-21] Overall 179 patients (18.8%) were carriers of at least one minor allele. All three SNPs were significantly associated with Lp(a) levels in the LipidCardio cohort (Table 3). The strongest evidence of an association with Lp(a) serum levels was found for rs10455872 (β=0.794; p= 5.1×10^-52^, Table 3 and Figure 1). Our data were compatible with an overall positive association (OR 1.35; 95 % CI 0.93-1.99, p=0.118) between carrying a minor *LPA* variant and CAD; even though the evidence was week. However, when stratified by sex there was reasonable evidence to suggest an association between the *LPA*-SNPs tested and obstructive CAD in women (OR 1.85, 95%CI 1.02-3.35, p=0.04), while in men there was no evidence for such an association. The association in women was stable even after adjusting for age, type 2 diabetes, LDL-C, and statin therapy.

**Table 3:** Associations of *LPA*-SNPs and Lp(a) serum levels: linear regression analysis on Lp(a) serum levels (log10-transformed), adjusted for age and sex

| SNP | Location<br>(GRCh37.p13) | A1/A2<br>(call rate, %) | MAF | N<sup>1</sup> | β | SE | p-value | N<sup>2</sup> |
| --- | --- | --- | --- | --- | --- | --- | --- | --- |
| rs10455872 | 6:160589086 | A/G (99.6) | 0.0736 | 950 | 0.794 | 0.049 | 5.10×10<sup>-52</sup> | 933 |
| rs3798220 | 6:160540105 | C/T(99.9) | 0.0236 | 953 | 0.865 | 0.092 | 3.93×10<sup>-20</sup> | 936 |
| rs186696265 | 6:160690668 | C/T (99.8) | 0.0170 | 952 | 0.993 | 0.106 | 7.60×10<sup>-20</sup> | 935 |
| rs429358 | 19:44908684 | C/T(95.4) | 0.1307 | 910 | 0.035 | 0.043 | 0.407 | 893 |
| rs7412 | 19:44908822 | C/T(99.9) | 0.0692 | 953 | 0.150 | 0.057 | 0.008 | 936 |

Notes: A1/A2= allele 1/ allele2; MAF = allele frequency of the minor allele in
LipidCardio; SE = standard error; N<sup>1</sup>=number of genotyped samples; N<sup>2</sup>=number analyzed with respect to Lp(a) association = genotype and Lp(a) level available

**Figure 1.**
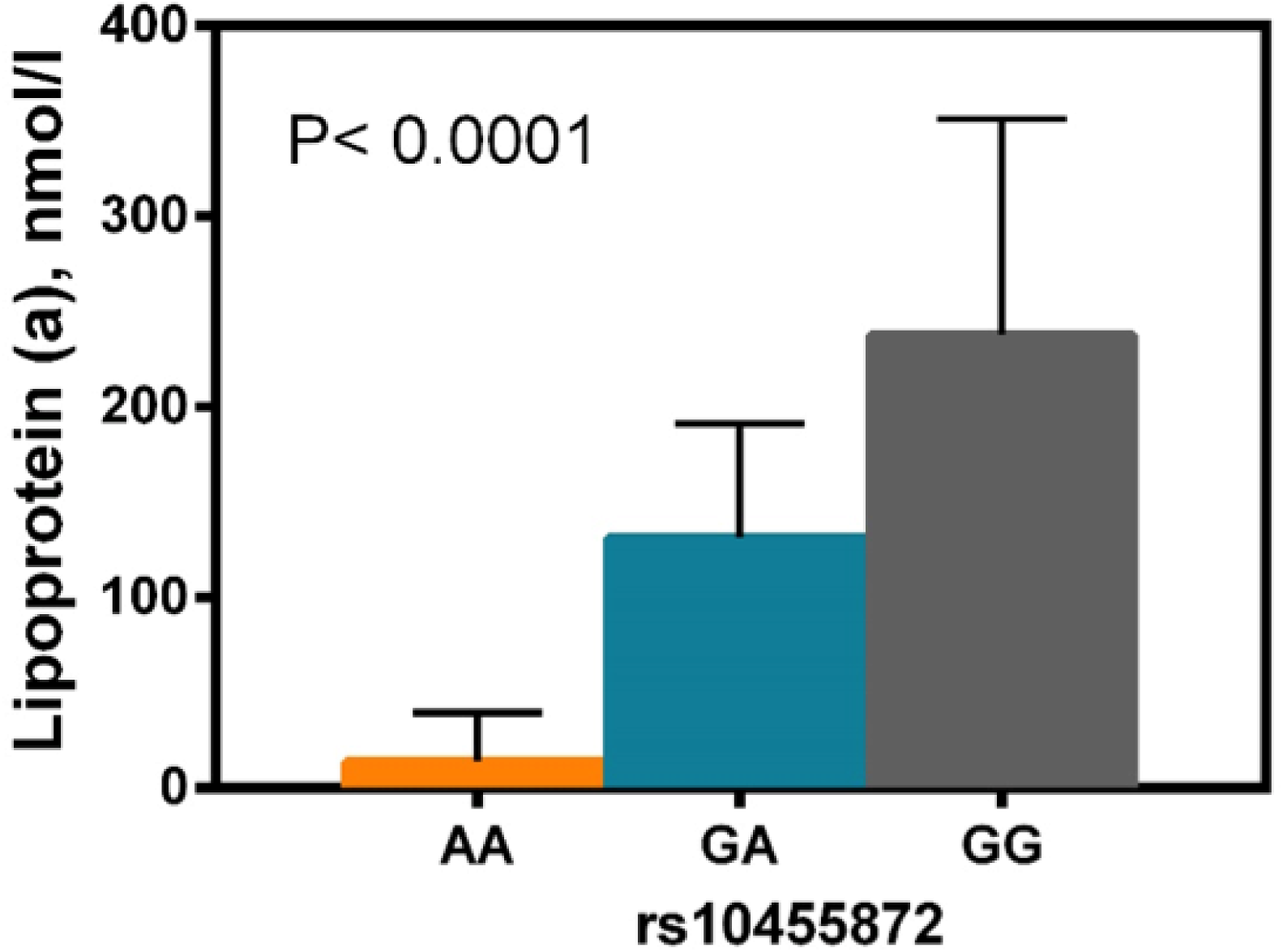
Association between SNP rs10455872 and Lipoprotein (a) in the LipidCardio sample. Rs10455872 genotypes were significantly associated with median Lp(a) values (interquartile ranges are indicated, N=933). The p-value was determined using the Kruskal-Wallis test.

Furthermore, we genotyped the two SNPs rs429358 and rs7412 constituting the three APOE isoforms apoE2, apoE3, and apoE4. Rs7412 was recently confirmed in a large GWAS (meta-analysis) to be associated with Lp(a) serum levels with genome wide significance. [18] Indeed, we found rs7412 to be significantly associated with Lp(a) concentration (β=0.15; p= 0.008), i.e. we could replicate the previous finding. Similar to the earlier study, there was no evidence that rs429358 was also associated with Lp(a) serum levels in the LipidCardio cohort (data not shown), while there was reasonable evidence of an association between the inferred APOE isoforms (**Supplementary Table 2**) and Lp(a) serum levels (β=0.015; p= 0.006, N=893) (see **Supplementary Table 2**).

### Strengths and limitations of this study

- Most importantly, this cohort has coronary angiographic information on all included subjects. Thus, while the majority has confirmed obstructive CAD, we can ensure that the unaffected “comparison group” effectively has no apparent CAD.
- In addition we are able to differentiate a group with non-obstructive CAD, constituting an interesting intermediate group. This differentiation is particularly important in terms of distinct phenotypes.
- A potential weakness is that patients are only recruited from one tertiary care hospital. Therefore, the results may not be generalizable to other patients with CAD.

## Supporting information

Supplementary meterial

## Collaboration

We are interested to share data and biobank samples with other researchers in joint collaborative projects, e.g. for replication of phenotypic and/or genetic findings or meta-analyzes of study results. Interested groups should contact the study coordinating PI Ilja Demuth at for the data-sharing application form. Each application will be reviewed by the LipidCardio PIs (currently I.D., U.L. and E.S-T.) and the decision communicated to the applicants usually within 6 weeks of submission.

## Funding

The LipidCardio Study was partially funded by the Sanofi-Aventis Deutschland GmbH. This funder did not play a role in the study design, data collection and analysis, decision to publish, or preparation of the manuscript and only provided financial support.

## Acknowledgements

We would like to thank Sanofi-Aventis Deutschland GmbH for financial support and I.E.M GmbH, Germany for support of the pulse wave analysis. We kindly acknowledge the excellent cooperation with the Central Biomaterial Bank, the joint core facility of the Charité-Universitätsmedizin Berlin and the Berlin Institute for Health.

We thank our student research assistants Andrea Schwarz, Fabiola Lugano, Marc Martinovic, Sabrina Bäther, Radostina Misirkova, and Ilona Enarovic for their excellent assistance during study implementation. Furthermore, we appreciate the great support by the whole team of the interventional cardiology unit.

## Contributor statement

Conceived and designed the study: ID, MK, UL, and EST. Recruitment of participants: UL, DL, MK and DS. Providing routine clinical data: UL and DL. Collected study specific data: MK, SJ, AS, DS, and ID. Analyzed the data: MK and ID. Wrote the manuscript: MK and ID. All authors revised and approved the manuscript.

