## Supplementary meterial for "Cohort profile: The LipidCardio Study - Role of Lipoproteins in Cardiovascular Disease"

König et al.

**Supplementary Table 1.** Summary of items available in the LipidCardio Study

| **Item** | **N** | **Source** | **Method/Commentary** |
|---|---|---|---|
| Sex, Age, Ethnic group | 1,005 | Face-to-face interview | n.a. |
| Previous diagnoses/personal history/allergies | 1,005 | Electronic medical record | more than 200 items |
| Coronary angiography information | 1,005 | coronary angiography | |
| Sodium (mmol/l) | 956 | Laboratory analysis | see http://www.laborberlin.com/leistungsverzeichnis |
| Potassium (mmol/l) | 962 | | |
| Calcium(mmol/l) | 207 | | |
| LDL-C (mg/dl) | 964 | | |
| HDL-C (mg/dl) | 961 | | |
| TC (mg/dl) | 731 | | |
| HbA1c (%) | 983 | | |
| Creatinine (mg/dl) | 972 | | |
| ASAT (U/L) | 707 | | |
| ALAT (U/L) | 715 | | |
| gGT (U/L) | 215 | | |
| aPTT (sec.) | 806 | | |
| INR | 945 | | |

| | | | |
|---|---|---|---|
| CRP (mg/L) | 705 | | |
| Glucose (mg/dl) | 753 | | |
| Lp(a) (nmol/L) | 968 | | |
| ApoB (g/L) | 956 | | |
| ApoA1 (g/L) | 955 | | |
| TSH (mU/L) | 906 | | |
| Troponin T hs (ng/L) | 380 | | |
| CK (U/L) | 737 | | |
| CK-MB (U/L) | 676 | | |
| NT-proBNP (ng/L) | 223 | | |
| Hemoglobin (g/dL) | 972 | | |
| Platelets (/nl) | 984 | | |
| Leukocytes (/nl) | 984 | | |
| DNA | 948 | selected SNPs have already been genotyped | |
| Leukocyte telomere length | >800 | will be soon available | |
| DNA methylation age | >800 | will be soon available | |
| Ejection fraction (%) | 805 | Echocardiography | Echocardiography |
| Septal diameter, diastolic (mm) | 761 | Echocardiography | Echocardiography |
| Diastolic dysfunction (yes/no) | 787 | Echocardiography | Echocardiography |
| Aortic valve stenosis (yes/no) | 821 | Echocardiography/Electronic medical record | Echocardiography |

| | | | |
|---|---|---|---|
| Medication (prescription and non-prescription drugs, including daily doses) | 1005 | Electronic medical record | |
| Height (cm) | 914 | self-reported, face-to-face interview | |
| Weight (kg) | 913 | self-reported, face-to-face interview | |
| BMI (kg/$m^2$) | 913 | calculated | |
| Waist circumference (cm) | 852 | clinical examination | measured in light clothes and standing position |
| Hip circumference (cm) | 851 | clinical examination | measured in light clothes and standing position |
| PWA (Pulse wave velocity, Augmentation @75 Index (AIx@75), Stroke Volume, Cardiac Output, Vascular Resistance) | 730 | clinical examination | Mobil-O-Graph® |
| Blood pressure (mm Hg) | 908 | clinical examination | BOSO medicus uno®<br>2 measurements (right arm, left arm) |
| Hand grip strength (kg) | 884 | clinical examination | Smedley Dynamometer (Saehan, Type SH5002); 3 measurements on both arms each |

| | | |
|---|---|---|
| Smoking (Start date, Stop date, Pack years) | 900 | Face-to-face interview |
| Alcohol | 888 | Face-to-face interview |
| Familiy health history including family tree | 896 | Face-to-face interview |
| MMSE | 830 | Face-to-face interview |
| Food preferences questionnaire | 807 | Questionnaire |
| RAPA | 847 | Questionnaire |
| Seattle Angina Questionnaire | 697 | Questionnaire |

Notes: Fluctuating of numbers is due to the fact that parts of the data were taken from the clinical routine if they were available, and logistical reason (e.g. questionnaires).

**Supplementary Table 2:** *APOE* genotypes and the resulting isoforms (N=893).

| rs7412 | rs429358 | APOE isoforms | N (%) |
|---|---|---|---|
| TT | TT | E2/E2 | 5 (0.5) |
| CT | TT | E2/E3 | 104 (11.4) |
| CT | CT | E2/E4 | 12 (1.3) |
| CC | TT | E3/E3 | 574 (63.1) |
| CC | CT | E3/E4 | 193 (21.2) |
| CC | CC | E4/E4 | 22 (2.4) |
